# LOMAR: LOcalization Microscopy Analysis in R

**DOI:** 10.1101/2022.05.30.493957

**Authors:** Maria Theiss, Alvis Brazma, Virginie Uhlmann, Jean-Karim Hériché

**Author notes:** Corresponding author (JKH).

## Abstract

In single molecule localization microscopy data (SMLM), individual instances of a macromolecular complex come in the form of point sets. Particle averaging, which combines localization data from a large number of instances, is often used to overcome experimental noise and obtain a refined view of the underlying structure. However, SMLM point sets are often heterogeneous due to biological variations in the structure they represent and must be partitioned into groups with similar structure before averaging which calls for being able to compute structurally-relevant similarity measures between sets of points. Here we introduce LOMAR (LOcalization Microscopy Analysis in R) a software package for the R programming language that enables comparison of point sets through implementation of several point sets registration methods and similarity measures derived from topological data analysis. We demonstrate use of the package on real and simulated SMLM data of nuclear pore complexes.

## Introduction

SMLM techniques enable the determination of a fluorophore’s position with a resolution of a few nanometers and is increasingly applied to the characterization of macromolecular complexes inside cells such as the nuclear pore complex (Sabinina et al., 2021), DNA damage repair foci (Sisario et al, 2018) or endocytic structures (Mund et al., 2018). While more typical light microscopy techniques produce data in the form of arrays of intensity values (i.e. images), SMLM produces sets of coordinates (i.e. point sets) with some additional information such as channel, precision and number of detected photons. While such data can be converted to images and processed using standard image analysis methods, new software and methods are have been developed to deal directly with the point cloud nature of SMLM data. Most of these focus on the segmentation of individual instances of the macromolecular structure of interest by applying clustering techniques to the coordinates data (Wu et al., 2020).

Incomplete labelling of structures and the stochastic nature of SMLM data acquisition result in noisy point sets with missing data. To overcome this issue, combining large numbers of instances with particle averaging (also sometimes called particle fusion) methods is increasingly used to refine the characterization of underlying structures. Particle averaging assumes that combined instances represent the same compositional and conformational structure but this assumption is not satisfied when biological variations contribute to the heterogeneity of the point sets obtained by SMLM. Partitioning point sets into structurally homogeneous groups is therefore necessary before applying particle averaging. This however requires being able to evaluate structural similarity between point sets in a manner that is robust to noise and missing data.

To help with this challenge, we developed the LOMAR software package which focuses on 3D point sets comparison by implementing point set registration methods and similarity measures based on topological data analysis. LOMAR is written in the R programming language. We demonstrate use of LOMAR on the registration of real 3D SMLM nuclear pore data and on the task of clustering simulated 3D instances of nuclear pores.

## Results

### Data input

LOMAR can ingest SMLM 2D and 3D point set data from character-delimited text files. Various filters can be applied to select a subset of the points and selected points can be further grouped using either cluster membership information included in the input file or by applying the DBSCAN clustering algorithm. Point sets can also be read from a series of images in TIFF format for use for example when corresponding objects have been obtained by image segmentation methods. For visualisation, and sometimes processing, it can be advantageous to convert point setss to images. For this, LOMAR implements a histogram binning method and an estimated photon count method (Huang et al., 2011).

### Point set registration methods

The objective of point set registration methods is to find correspondence between multiple point sets and identify the spatial transformation that optimally aligns them. These methods have a long history in fields where sensors are in use such as robotics (Pomerleau et al., 2015). However, many such classical methods are implemented in different and often non-interoperable specialised software which may limit their adoption for use with SMLM data. LOMAR brings together a few of these methods into one package (table 1): iterative closest point (ICP, Besl and McKay, 1992), coherent point drift (CPD, Myronenko and Song, 2010), and joint registration of multiple point clouds (JRMPC, Evangelidis et al., 2014) and a new pairwise registration of Gaussian mixture models using the Wasserstein distance (WGMMreg). Only rigid registration (i.e. involving only rotation and translation) is implemented for each method because structures produced by deformations should be considered as separate from the structures they can be derived from and detecting such structural variations is of biological relevance.

**Table 1:** Point set registration methods available in LOMAR.

| Method name | Acronym | Characteristics | Reference |
| --- | --- | --- | --- |
| Iterative closest point | ICP | • Processes a pair of point sets<br>• Based on distances between matching points<br>• Fast<br>• Sensitive to noise, outliers and missing points<br>• Converges to a local minimum | Besl and McKay, 1992 |
|  |  | Doesn't make use of additional SMLM information |  |
| Coherent point drift | CPD | Processes a pair of point sets<br>Maximises likelihood of a GMM<br>More robust to noise, outliers and missing points<br>Compromise between accuracy and computational complexity<br>May not be suitable for anisotropic SMLM data | Myronenko and Song, 2010 |
| Registration of Gaussian mixture models with Wasserstein distance | WGMMreg | Processes pair of point sets<br>Maximises likelihood of a GMM<br>Can use most available SMLM information<br>Slow due to high computational complexity | This paper |
| Joint registration of multiple point clouds | JRMPC | Simultaneously processes multiple point sets<br>Considers point sets as generated by a GMM + transformations<br>Robust to noise, outliers and missing points | Evangelidis et al., 2014 |

ICP is one of the most used registration methods due to its simplicity. The algorithm is a pairwise method that iterates between two steps: 1- given a rigid transformation (translation and rotation), assign correspondence between two point sets A and B by finding the closest points in A for all points in B, 2- given the points correspondence, find the best rigid transformation that aligns B with A. Although ICP is sensitive to starting conditions and noise, it is fast and often provides good enough alignments. However, better results are often obtained with methods involving Gaussian mixture models (GMMs) which allow incorporation of additional information such as level of noise or point localisation uncertainty.

Among these, CPD is a widely used pairwise registration method. In CPD, a Gaussian mixture model (GMM) is constructed from point set A and points in set B are considered observations from this GMM. The GMM centroids are then moved to maximise the likelihood of point set B. A coherence constraint is imposed on the movement to preserve the structure of the point sets. Fuzzy point set correspondences and the inclusion of a noise factor contribute to CPD’s relative robustness to noise and outliers. Additional information can be included in the form of a points correspondence weight matrix. However, to ensure convergence, the algorithm uses equal isotropic covariances which precludes incorporating localization precision in the covariances if it is anisotropic.

To remedy this situation, we developed WGMMreg, an algorithm to register two GMMs by minimising the Wasserstein distance between them. An algorithm called GMMreg has already been described (Jian and Vemuri, 2011) and is based on the minimization of the L2 distance between GMMs. However, use of the L2 distance discards information about the relative positions of the points. A natural alternative to compare probability distributions is the Wasserstein distance. It also has the advantage of including information about the relative positions of the points due to the computation of an optimal transport map. However, while the Wasserstein distance can be computed in closed form between two Gaussian distributions, the Wasserstein distance between GMMs (linear combinations of Gaussians) is intractable. In our WGMMreg implementation we leverage recent results on the computation of a discrete form of a Wasserstein-type distance between GMMs (Delon and Desolneux, 2020).

The computational cost of the registration methods described above increases with the amount of information included and in particular, our current implementation of WGMMreg becomes prohibitive for large point sets although this can be mitigated with downsampling. In addition, when registering a large number of point sets, the above methods require computing registrations between all pairs of points and subsequently using iterative methods (for example similar to the Barton-Sternberg algorithm for multiple sequence alignment (Barton and Sternberg, 1987)) to obtain a global registration map (see e.g. Heydarian et al., 2021). This also clearly becomes computationally expensive as the number of point sets increases.

For increased performance when many point sets need to be registered, we also implemented a method for the joint registration of multiple point clouds (JRMPC, Evangelidis et al., 2014). In this approach, the point sets are considered generated by a common GMM and both the mixture and the registration parameters are estimated via an expectation maximisation (EM) algorithm. In addition to the registered point sets, the algorithm returns the transformation associated with each point set and the final GMM. A form of model selection can also be applied by removing unsupported components of the GMM using a minimum message length approach (Figueiredo and Jain, 2002). To illustrate use of this method, we applied it to the registration of 356 NUP107 nuclear pores obtained by STORM (Heydarian et al, 2021). The JRMPC method produces a global registration revealing the 8 subunits structure (figure 1a,b) in less than 20 min using only one thread on a 3.6 GHz CPU. In contrast, Heydarian et al. report that the all-vs-all registration of the same data takes 2 h using parallelization on a GPU. With JRMPC, the resulting GMM can also provide information about the underlying structure and potential outliers can be detected from the high variance of the components they are assigned to. In the case of NUP107 whose primary structure consists of two rings, this identifies the between-rings components as outliers with the other components forming a double ring structure (figure 1b). Eliminating points associated with the high variance components improves the overall registration outcome (figure 1a, compare left and right panels). The joint registration of multiple point clouds can also be used to quickly produce a good initial state for iterative methods leading to potential gains in computation time.

**Figure 1:**
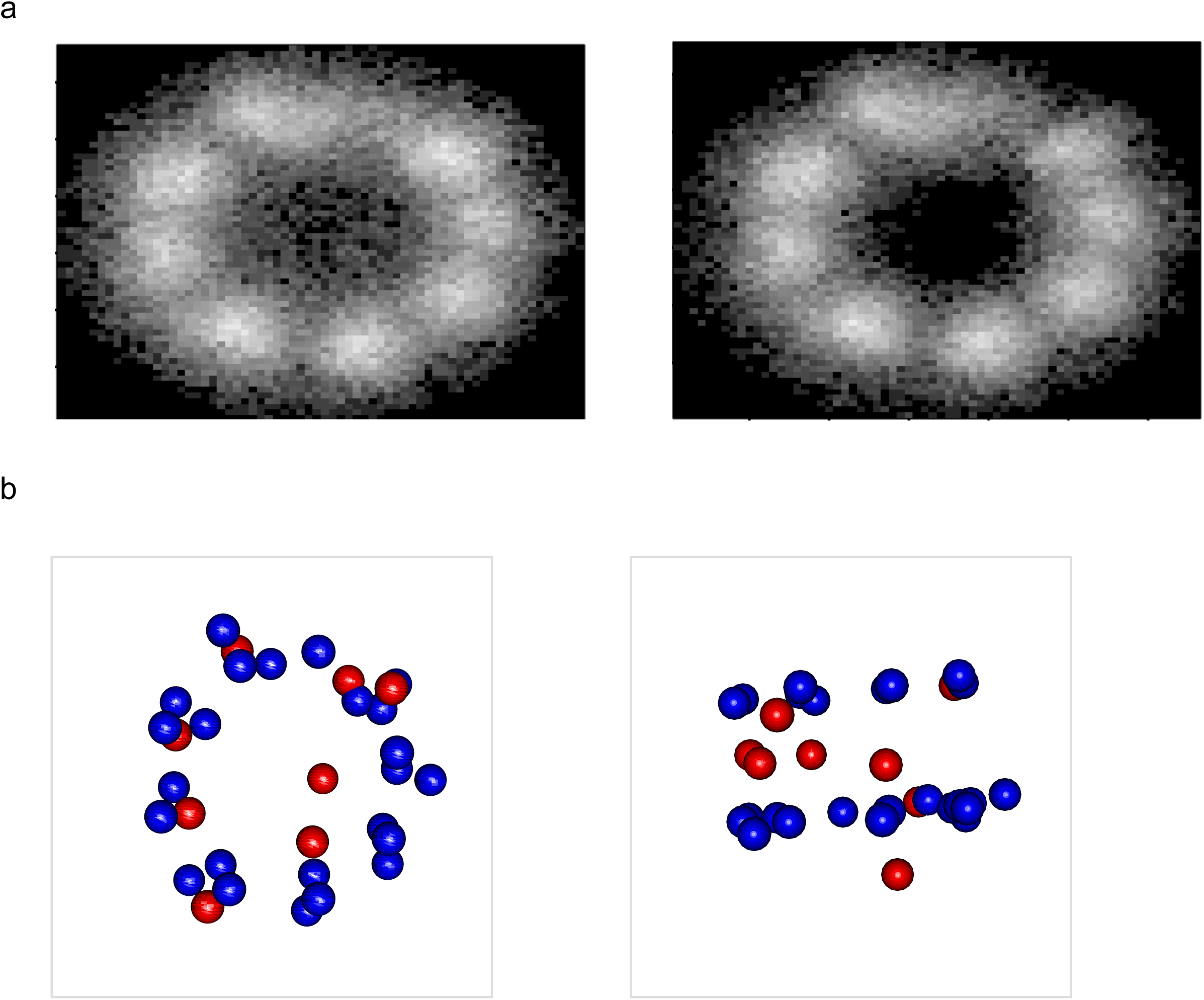
Joint registration of NUP107 point sets a) Joint registration of the 356 NUP107 point sets from Heydarian et al. rendered as an image using the histogram method. Left panel: without denoising, right panel: after denoising by removal of points associated with high variance components of the model. b) Model of NUP107 obtained by joint registration of all point sets. High variance components are shown in red.

### Topological data analysis

In topological data analysis (TDA), a set of data points with a measure of distance between them is seen as representing an underlying topological space. Relevant topological characteristics can be extracted from such topological spaces using the tools of persistent homology (Edelsbrunner and Harer, 2010; Otter et al, 2017, Amézquita et al, 2020). Informally, homology counts the number of “holes’’ of different dimensions in a topological space, i.e. the 0th homology group counts the number of connected components, the first homology group counts the number of (2d) holes (i.e. holes inside loops), the 2nd homology group counts the number of (3d) voids (i.e. bubbles). Persistent homology efficiently computes homology groups at multiple scales. For this, each point set is converted into a simplicial complex based on a proximity parameter t. A simplex is an n-dimensional triangle and a simplicial complex is a collection of simplices connected by following specific rules to ensure that the homology of the underlying topological space is preserved (figure 2a). Recording when holes form and get filled over a range of values of t produces a persistence diagram (figure 2b). To compute simplicial complexes and persistence diagrams, LOMAR relies on the Dionysus library with code borrowed from the TDA package (Fasy et al., 2014). With this approach, each point set of a SMLM data set can be represented by a persistence diagram that characterises the shape of its underlying structure. Because persistent homology relies only on the pairwise distance between points, the shapes captured by persistence diagrams are independent of position and coordinate system. Persistent homology is robust to noise in the sense that small perturbations to the points positions will minimally affect the persistence diagram. Outliers are of more concern since they can have a large effect on the persistence diagram. However, as they tend to be further away from other data points they can either be detected and removed beforehand or various techniques can be used to deal with this issue such as de-emphasizing low density regions, for example by the use of a distance-to-measure function (Chazal et al., 2018).

**Figure 2:**
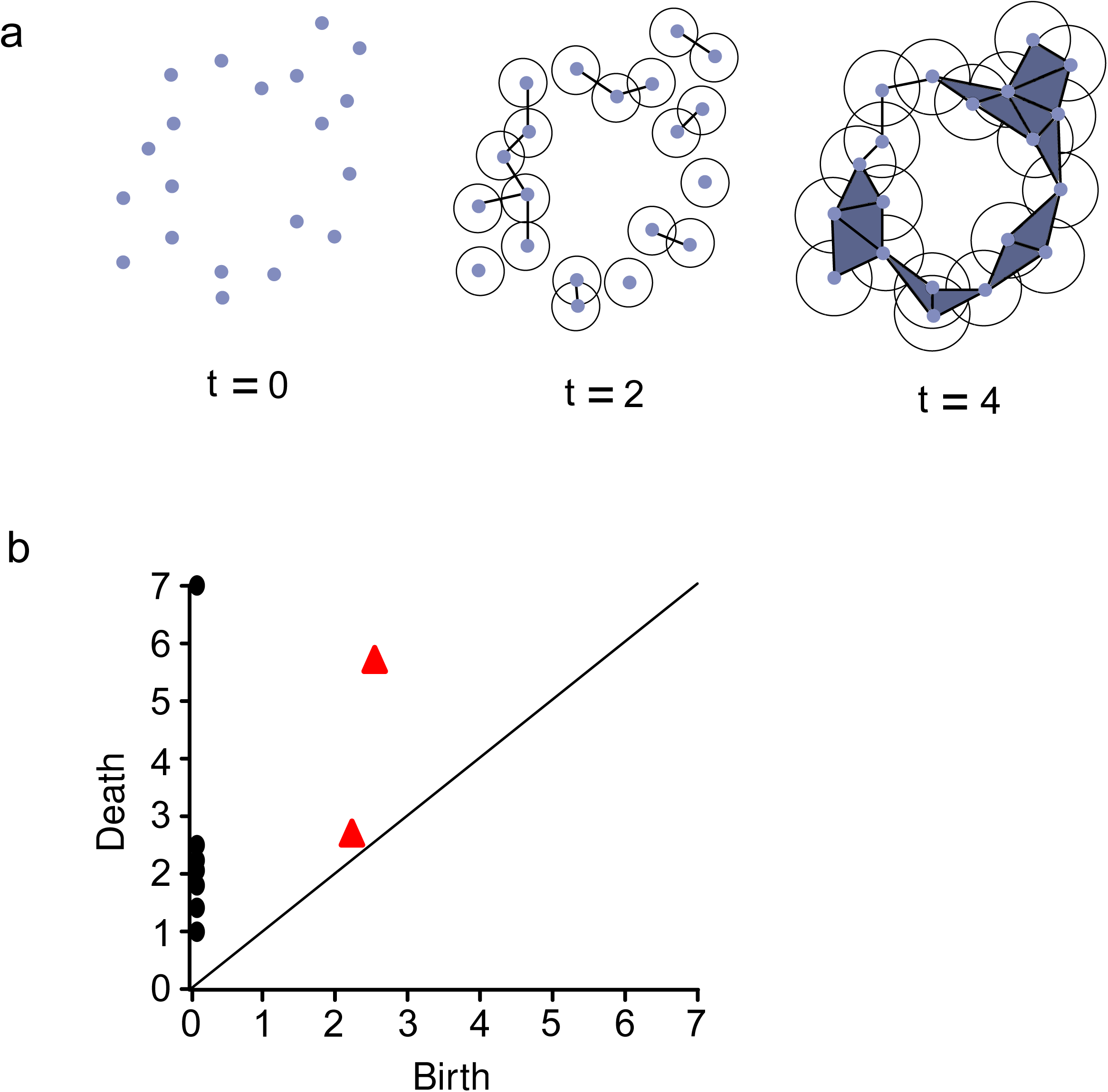
Persistent homology. a) Building a simplicial complex out of data points. The Vietoris-Rips complex for parameter t is built by including an edge between two points if they are within distance t of each other (i.e. the circles of diameter t centred on each point overlap). A higher dimensional simplex is included if all its possible edges are present, i.e. all the points that are members of the simplex are within distance t of each other. Here only 2-simplices (colored triangles) are shown. b) Persistence diagram of the point set shown in (a). Dots represent the 0D homology group (connected components) and triangles represent the 1D homology group (holes). Birth and death values are the values of t at which a feature respectively is formed and disappears.

### Similarity measures for point sets

There are several ways of deriving structurally-meaningful similarity between two sets of points with LOMAR. One is to make use of registration errors obtained from the application of registration algorithms such as those described above. Another is to view the point sets as discrete samples from probability distributions and use a suitable measure of similarity between probability distributions such as the Wasserstein distance. Point sets shape similarity can also be evaluated by comparing their persistence diagrams. Conceptually, the first two approaches capture shape similarity by evaluating how much displacement is required to align the point sets while the TDA approach could be seen as comparing the multi-scale “porosity” of the underlying structures. These approaches are therefore complementary. For compatibility with many machine learning methods which require positive definite similarity matrices, LOMAR implements the sliced Wasserstein distance between persistence diagrams (Carrière et al., 2017) and the persistence scale-space kernel (Reininghaus et al, 2015). The sliced Wasserstein distance is an approximation of the Wasserstein distance that is efficiently computed as a sum of Wasserstein distances between 1D projections of the points and produces a negative definite distance matrix. The persistence scale-space kernel results from mapping the persistence diagrams to a suitable vector space and computing a heat diffusion kernel.

### Clustering point sets

To illustrate the use of similarity measures for clustering SMLM point sets, we use simulated data inspired by the nuclear pore complex. Ground truth data is produced using the method from (Theiss et al. 2022) and its acquisition in a typical SMLM experiment is simulated using the SMAP software (Ries, 2020).

For TDA-based clustering, a persistence diagram of sublevel sets of the distance to measure function is computed for each point set then the matrix of (sliced) Wasserstein distances between persistence diagrams is computed. For registration-based clustering, the (symmetrized) matrix of registration errors obtained from the pairwise application of the cpd algorithm is used as a distance matrix.

Following (Huijben et al. 2021), the distance matrix is used to embed the point sets into a low dimension space using multidimensional scaling. Clusters are then recovered by Gaussian mixture modelling of the data points in this space. To evaluate clustering quality, we assign to each cluster the label of the class most represented in that cluster and compare assigned labels to the ground truth class of origin.

NPCs tagged on different subunits produce structures with different ring diameters or different distances between the two rings (Sabinina et al., 2021). To illustrate how such structures can be separated, we produced 3 classes of simulated NPC structures that differ by their average ring diameter or by the average distance between the two rings (figure 3a).

**Figure 3:**
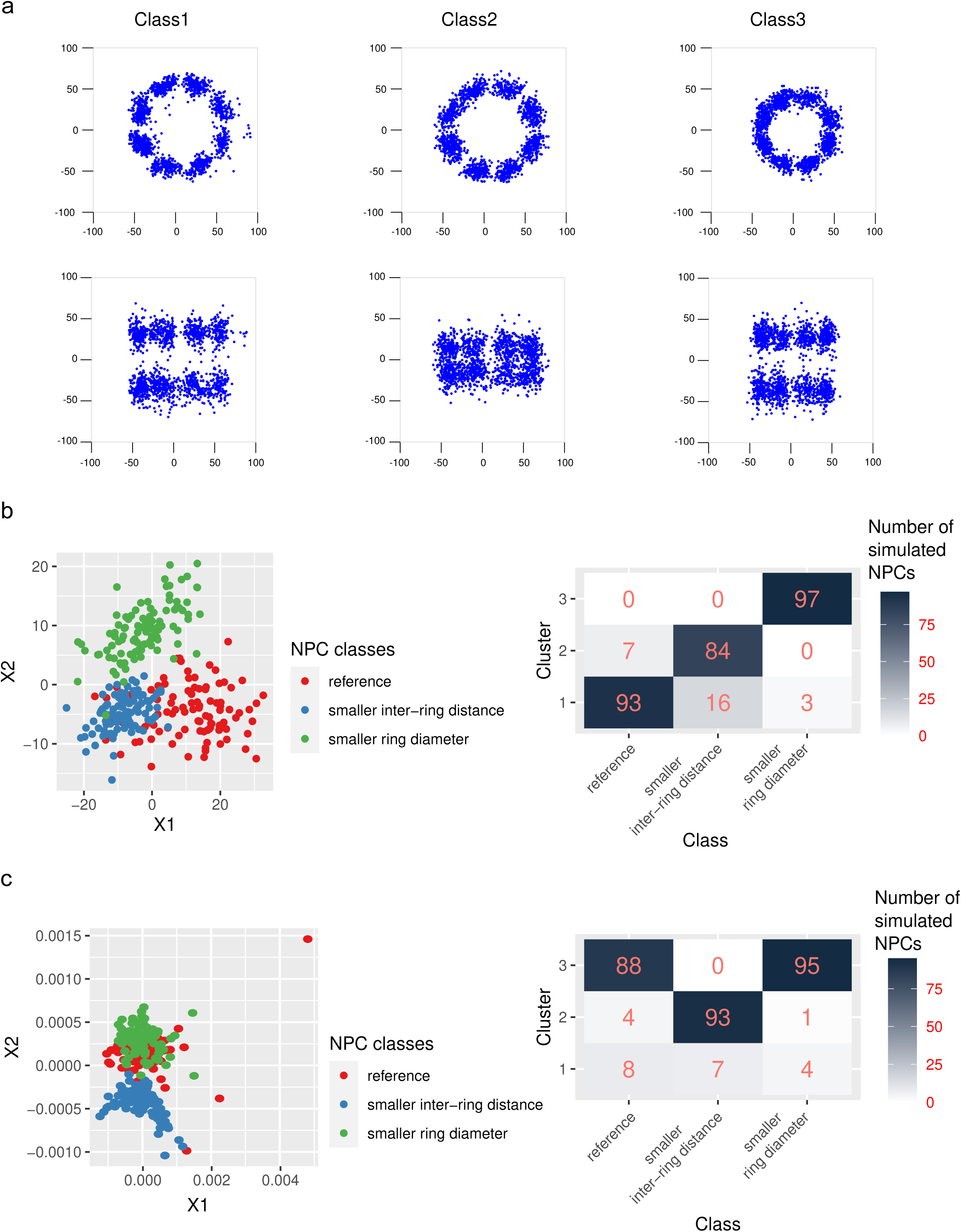
Clustering of multiple classes of simulated nuclear pore complexes a) Registered examples of point sets from 3 classes. Top row: top view, bottom row: side view. Class 1: reference class, average ring diameter: 60 nm, average inter-ring distance: 60 nm, class 2 : average ring diameter: 60 nm, average inter-ring distance: 30 nm, class 3: average ring diameter: 38.4 nm, average inter-ring distance: 60 nm. b) Clustering using TDA. Left panel: Points sets embedded in the first two dimensions of the multidimensional scaling feature space. Colours indicate the classes. Right panel: cluster composition. c) Clustering using registration. Left panel: Points sets embedded in the first two dimensions of the multidimensional scaling feature space. Colours indicate the classes. Right panel: cluster composition.

Using TDA, the three classes can be separated with an average accuracy of 91% (figure 3b) while registration-based methods fail to separate the two classes that differ by their ring diameter (figure 3c).

NPCs typically have rings with an 8-fold radial symmetry but a small proportion may have a 9-fold symmetry (Hinshaw and Milligan, 2003). To demonstrate clustering of SMLM structures in the presence of class imbalance, we simulated an NPCs data set in which 15% of the structures have a 9-fold symmetry (figure 4a). Here the TDA-based approach succeeds with an accuracy of 99% (figure 4b) while registration-based clustering fails to separate the two classes (figure 4c).

**Figure 4:**
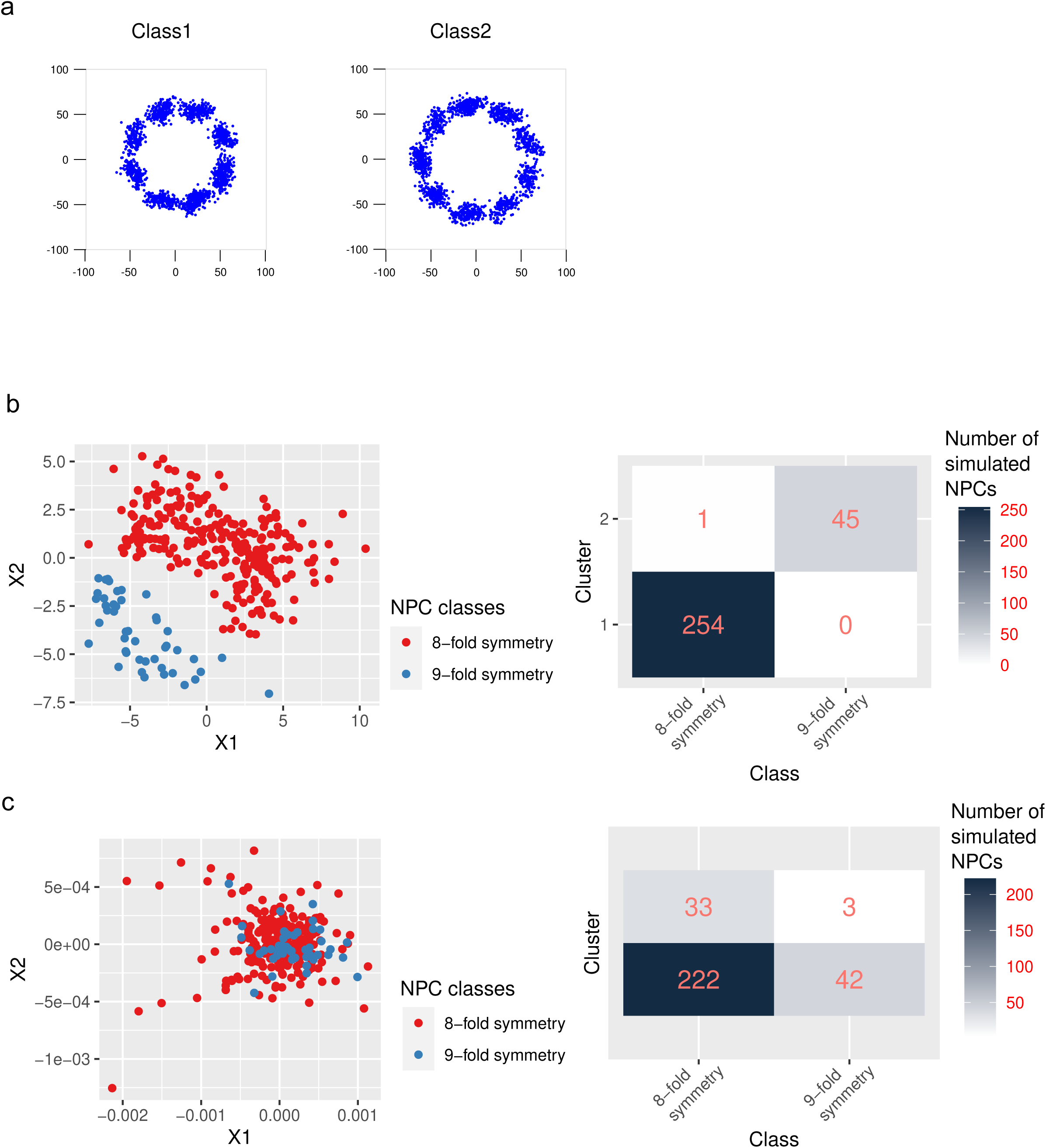
Clustering with class imbalance a) Registered examples of point sets from 2 classes. Class 1: rings with 8-fold symmetry, class 2 : rings with 9-fold symmetry. b) Clustering using TDA. Left panel: Points sets embedded in the multidimensional scaling feature space. Colours indicate the classes. Right panel: cluster composition. c) Clustering using registration. Left panel: Points sets embedded in the first two dimensions of the multidimensional scaling feature space. Colours indicate the classes. Right panel: cluster composition.

## Conclusion

We developed the LOMAR package to enable comparison of structures generated by single molecule localisation microscopy. To this end, LOMAR implements three sets of functionalities: reading of SMLM data, point set registration methods and persistent homology-based measures of similarity between point sets. While the motivation for its development was the analysis of 3D single molecule localization microscopy data, most functions are generic and can be applied to any 2D or 3D point sets.

## Availability

The LOMAR package is available on the R package repository CRAN: https://cran.r-project.org/package=LOMAR and the source code is available from a GitLab repository at https://git.embl.de/heriche/lomar. Data and code notebooks used to produce material for the figures are available at https://git.embl.de/heriche/lomar_use_examples.

## References

Sabinina VJ, Hossain MJ, Hériché JK, Hoess P, Nijmeijer B, Mosalaganti S, Kueblbeck M, Callegari A, Szymborska A, Beck M, Ries J, Ellenberg J. Three-dimensional superresolution fluorescence microscopy maps the variable molecular architecture of the nuclear pore complex. Mol Biol Cell. 2021 Aug 15;32(17):1523–1533. doi: 10.1091/mbc.E20-11-0728. Epub 2021 Jun 30. PMID: 34191541; PMCID: PMC8351745.

Sisario D, Memmel S, Doose S, Neubauer J, Zimmermann H, Flentje M, Djuzenova CS, Sauer M, Sukhorukov VL. Nanostructure of DNA repair foci revealed by superresolution microscopy. FASEB J. 2018 Jun 12:fj201701435. doi: 10.1096/fj.201701435. Epub ahead of print. PMID: 29894665.

Mund M, van der Beek JA, Deschamps J, Dmitrieff S, Hoess P, Monster JL, Picco A, Nédélec F, Kaksonen M, Ries J. Systematic Nanoscale Analysis of Endocytosis Links Efficient Vesicle Formation to Patterned Actin Nucleation. Cell. 2018 Aug 9;174(4):884-896.e17. doi: 10.1016/j.cell.2018.06.032. Epub 2018 Jul 26. PMID: 30057119; PMCID: PMC6086932.

Wu YL, Tschanz A, Krupnik L, Ries J. Quantitative Data Analysis in Single-Molecule Localization Microscopy. Trends Cell Biol. 2020 Nov;30(11):837–851. doi: 10.1016/j.tcb.2020.07.005. Epub 2020 Aug 20. PMID: 32830013.

Huang F, Schwartz SL, Byars JM, Lidke KA. Simultaneous multiple-emitter fitting for single molecule super-resolution imaging. Biomed Opt Express. 2011 Apr 29;2(5):1377–93. doi: 10.1364/BOE.2.001377. PMID: 21559149; PMCID: PMC3087594.

Pomerleau F, Colas F, Siegwart R. A Review of Point Cloud Registration Algorithms for Mobile Robotics. Foundations and Trends in Robotics, Now Publishers, 2015, 4 (1), pp.1--104. (10.1561/2300000035) (hal-01178661)

Besl P, McKay. H A method for registration of 3-D shapes. Pattern Analysis and Machine Intelligence, IEEE Transactions on, 14(2):239–256, February 1992.

Myronenko A, Song X. Point set registration: coherent point drift. IEEE Trans Pattern Anal Mach Intell. 2010 Dec;32(12):2262–75. doi: 10.1109/TPAMI.2010.46. PMID: 20975122.

Evangelidis G, Kounades-Bastian D, Horaud R, Psarakis E. A Generative Model for the Joint Registration of Multiple Point Sets. European Conference on Computer Vision, Sep 2014, Zurich, Switzerland. pp.109–122, (10.1007/978-3-319-10584-0_8). (hal-01019661v3)

Jian B, Vemuri BC. Robust Point Set Registration Using Gaussian Mixture Models. IEEE Trans Pattern Anal Mach Intell. 2011 Aug;33(8):1633–45. doi: 10.1109/TPAMI.2010.223. Epub 2010 Dec 23. PMID: 21173443.

Delon J, Desolneux A. A Wasserstein-Type Distance in the Space of Gaussian Mixture Models. SIAM J. Imaging Sci. 13 (2020): 936–970.

Barton GJ, Sternberg MJ. A strategy for the rapid multiple alignment of protein sequences. Confidence levels from tertiary structure comparisons. J Mol Biol. 1987 Nov 20;198(2):327–37. doi: 10.1016/0022-2836(87)90316-0. PMID: 3430611.

Heydarian H, Joosten M, Przybylski A, Schueder F, Jungmann R, Werkhoven BV, Keller-Findeisen J, Ries J, Stallinga S, Bates M, Rieger B. 3D particle averaging and detection of macromolecular symmetry in localization microscopy. Nat Commun. 2021 May 14;12(1):2847. doi: 10.1038/s41467-021-22006-5. Erratum in: Nat Commun. 2021 May 26;12(1):3262. PMID: 33990554; PMCID: PMC8121824.

Figueiredo MAT, Jain AK. Unsupervised learning of finite mixture models. IEEE Transactions on Pattern Analysis and Machine Intelligence, vol. 24, no. 3, pp. 381–396, March 2002, doi: 10.1109/34.990138.

Edelsbrunner H, Harer J. Computational Topology: An Introduction. American Mathematical Society. 2010. 10.1007/978-3-540-33259-6_7.

Otter N, Porter MA, Tillmann U, Grindrod P, Harrington HA. A roadmap for the computation of persistent homology. EPJ Data Sci. 2017;6(1):17. doi: 10.1140/epjds/s13688-017-0109-5. Epub 2017 Aug 9. PMID: 32025466; PMCID: PMC6979512.

Amézquita EJ, Quigley MY, Ophelders T, Munch E, Chitwood DH. The shape of things to come: Topological data analysis and biology, from molecules to organisms. Dev Dyn. 2020 Jul;249(7):816–833. doi: 10.1002/dvdy.175. Epub 2020 Apr 13. PMID: 32246730; PMCID: PMC7383827.

Fasy B, Kim J, Lecci F, Maria C. Introduction to the R Package TDA. 2014. arXiv preprint 1411.1830.

Chazal F, Fasy B, Lecci F, Michel B, Rinaldo A, Wasserman L. Robust Topological Inference: Distance To a Measure and Kernel Distance. Journal of Machine Learning Research, Microtome Publishing, 2018, 18 (159), pp.40. (hal-01232217)

Carrière M, Cuturi M, Oudot S. Sliced Wasserstein kernel for persistence diagrams. In Proceedings of the 34th International Conference on Machine Learning – 2017. Volume 70 (ICML’17). JMLR.org, 664–673.

Reininghaus J, Huber S, Bauer U, Kwitt R. A stable multi-scale kernel for topological machine learning. In Proceedings of the IEEE Conference on Computer Vision and Pattern Recognition(CVPR), pages 4741–4748, 2015.

Theiss M, Hériché JK, Russell C, Helekal D, Soppitt A, Ries J, Ellenberg J, Brazma A, Uhlmann V. Simulating structurally variable Nuclear Pore Complexes for Microscopy. bioRxiv 2022 May 17;492295.

Ries J. SMAP: a modular super-resolution microscopy analysis platform for SMLM data. Nat Methods. 2020 Sep;17(9):870–872. doi: 10.1038/s41592-020-0938-1. PMID: 32814874.

Huijben TAPM, Heydarian H, Auer A, Schueder F, Jungmann R, Stallinga S, Rieger B. Detecting structural heterogeneity in single-molecule localization microscopy data. Nat Commun. 2021 Jun 18;12(1):3791. doi: 10.1038/s41467-021-24106-8. PMID: 34145284; PMCID: PMC8213809.

Hinshaw JE, Milligan RA. Nuclear pore complexes exceeding eightfold rotational symmetry. J Struct Biol. 2003 Mar;141(3):259–68. doi: 10.1016/s1047-8477(02)00626-3. PMID: 12648571.

